# The geometry of admixture in population genetics: the blessing of dimensionality

**DOI:** 10.1101/2023.09.08.556908

**Authors:** José-Angel Oteo, Gonzalo Oteo-García

## Abstract

We present a geometry-based interpretation of the *f*-statistics framework, commonly used to determine phylogenetic relationships from genetic data. The focus is on the determination of the mixing coefficients in population admixture events subject to post-admixture drift. The interpretation takes advantage of the high dimension of the dataset and analyzes the problem as a dimensional reduction issue. We show that it is possible to think of the *f*-statistics technique as an implicit transformation of the genetic data from a phase space into a subspace where the mapped data structure is more similar to the ancestral admixture configuration. The positive effect of the map can be explicitly assessed. The overarching geometric framework provides slightly more general formulas than the *f*-formalism by using a different rationale as a starting point. Explicitly addressed are two- and three-way admixtures. The mixture proportions are provided by suitable linear fits in two or three dimensions that can be easily visualized. The developments and findings are illustrated with numerical simulations from real world datasets.

## 1 Introduction

The determination of admixture proportions in hybrid populations is a central topic in population genetics. At large evolutionary time scales, a population admixture event can be thought of as a sudden process in which a new linage emerges as the genetic weighted combination of two or more donors, in its simplest formulation.

The formalism known as *f*-statistics [1, 2, 3, 4, 5, 6, 7], which deals with allele-sharing correlations between two, three and four populations, is a commonly used technique to determine the admixture coefficients. Despite their simple computation and definition, the *f*-statistics outcome assessment may not be straightforward [5]. We develop a geometry-based interpretation of *f*-statistics intended to deepen understanding of the population admixture problem. Geometric-like methods have already been raised in the past [8, 9, 10, 11, 12, 13]. The strength of our approach is to take advantage of the high dimensionality of the problem, broadening the *f*-formalism toolkit.

Whenever the genetic information from the original populations is available, determining the admixture proportions is straightforward. However, the evolutionary history of the original populations manifests itself, for example, in random changes in allele frequencies, a phenomenon known as genetic drift. Although ancient DNA can sometimes aid in estimating admixture proportions by providing better proxies, the time elapsed since admixture and the possibility that the putative parents themselves are extinct make accurate estimates difficult. Moreover, a population may experience over time continuous, discrete, occasional, or repeated genetic influx from other or multiple others, which makes the issue more challenging. The present study is concerned with the approximation of admixture proportions in populations that are assumed to have evolved in time via genetic drift. A dataset with population allele frequencies corresponding to a large number of Single Nucleotide Polymorphisms (SNPs) is the common starting point for dealing with this question.

The population admixture problem may be posed as a geometric issue in a phase space, the allele frequency space, where each population is represented by a point and each axis corresponds to a SNP in the dataset [14]. With only three SNP, Figure 1 outlines the nature of the problem. Populations *i, a*′, *b*′, *a, b, j*, are assumed to be related by phylogenetic treeness and are located in phase space by their position vectors whose coordinates are given by the allele frequencies in the dataset. At the time of admixture, we have three co-linear points, *a*′, *x*′, *b*′, with *a*′ and *b*′ the donors and *x*′ the hybrid.

**Figure 1:**
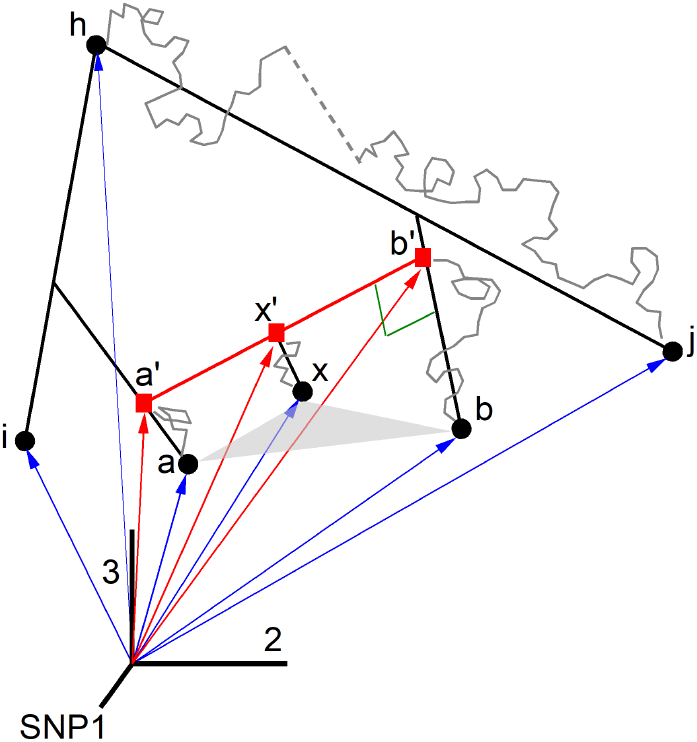
Three dimensional allele frequency space. Populations *a*′, *b*′ are the donors and *x*′ the hybrid at admixture birth. They evolve to a, x, b (proxies in the dataset) via genetic drift (Brownian-like gray curves). The Brownian-like trajectory from h to j is broken up by a random jump, which is intended to represent an event such as a population bottleneck. The triangle Δ axb is useful in the discussion because the geometric condition for *x* to be a hybrid of a and b is that the angle at vertex *x* be obtuse.

Then, they describe Brownian-like trajectories because allele frequencies undergo small random changes as a result of genetic drift, resulting in population proxies *a, x, b.*

Populations *i* and *j*, which appear as spectators in the Figure 1, will be referred to as auxiliary populations and will play an important role. Any population related to the admixture contributors via phylogenetic treeness may be considered as auxiliary. The Brownian-like trajectory depicted from *h* to *j* is interrupted by a *random jump*, which is meant to represent a sudden event that is prone to occur in long evolutionary histories. For instance, in a population bottleneck allele frequencies suffer a sudden change which translates in the random jump in phase space represented by the dashed line in the figure.

The geometric condition for *x* to be a hybrid is 90° < *ϕ* < 180°, where *ϕ* is the angle at vertex *x* in triangle Δ *axb*. Otherwise, whenever the three angles are acute, the populations *a, x, b*, are mathematically related by phylogenetic *treeness* [14]. A difficulty associated with long evolutionary histories is that an admixture triangle in phase space may evolve to have three acute angles, fooling the negative cosine test, cos *ϕ* < 0, as in Figure 2. And the way around, a treeness scenario might appear mathematically as admixture. This is why complementary knowledge about the population history is important.

**Figure 2:**
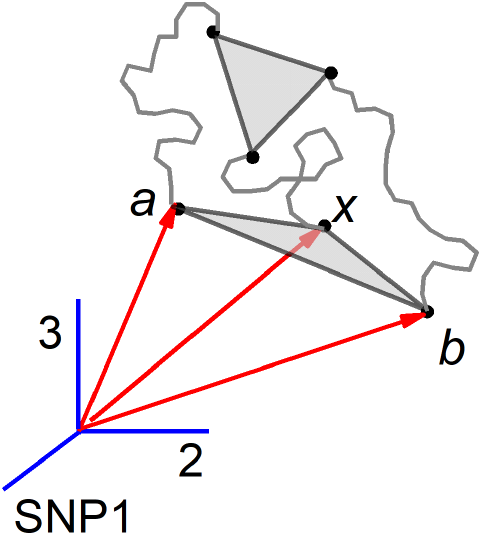
Fooling the admixture test. The parents a, b and the hybrid *x* form the triangle Δ axb, which is a post-admixture configuration that still evolves in time to a configuration corresponding to treeness because the obtuse angle at vertex *x* becomes acute. This description read in reverse corresponds to a treeness configuration becoming an apparent admixture.

The population admixture issue that we are addressing here is stated as follows. Determine the relative length of the segment 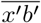 with respect to 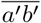, when we only know the triangle Δ *axb*. The solution provides an estimate of the admixture proportions. No reconstruction of the ancestral allele frequencies is contemplated.

A crucial ingredient in the approach is the high dimension of the allele frequency space, fixed by the number of SNP sequenced, typically in the hundreds of thousands. Geometry in high dimensions presents challenges because the space is *too empty*. The conventional example to illustrate it considers the volume of a cube of side 2 in ℝ^*n*^, which is 2^*n*^ bigger than the volume of a unit cube, despite the fact that their sides are only a factor 2 apart. *Curse of dimensionality* is the term used to characterize this feature. Fortunate enough, there are also positive derivatives associated to high dimensionality that will be helpful to approach the admixture problem. In this sense, we will use (i) the Johnson-Lindenstrauss lemma about dimension reduction, which states that pairwise distances are approximately preserved after projection onto a random subspace; (ii) the Brownian-like character (induced by genetic drift) of the population trajectories along the time in phase space, and (iii) the quasi-orthogonality of random vectors in high dimension.

In the next Section we describe in detail the overarching framework and its relationship with the standard *f*-statistics formalism. Section 3 gathers a number of simulations based on real world data intended to illustrate every aspect described in the preceding section. Section 4 contains final comments and results discussion. One 4 of technical character is included.

## 2 Population admixture, geometry and *f* -statistics

Consider the case of two populations thought to be the contributors to a third, hybrid, population. Assume that estimates of the allele frequencies of a number s of SNP are known. These three populations are the points *a, b, x,* respectively, in Figure 1, where s = 3. They are the proxies of the ancestral populations *a*′, *b*′, *x*′, extant when the admixture takes place. Indeed, the crinkly trajectories separating a, b, *x* from *a*′, *b*′, *x*′ are the knot of the population admixture problem. It is important to note that the diagram hosts a number of features that cannot be consistently represented in 3D. For instance, the points structure {*a*′, *x*′, *b*′, *b, x, a*} is by no means planar because vectors from *a*′ to a, from *x*′ to *x*, and from *b*′ to *b*, point towards random directions. Moreover, the genetic drift that drives population *b*′ to *b* is such that the segment 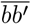 is approximately orthogonal to 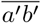, a fact that is indicated by the squared (green) angle. There exist further quasi-orthogonality (denoted ⊥) constraints: 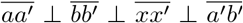. This quasiorthogonality-based description, which is justified bellow, is a consequence of the dynamics induced by genetic drift in a high dimension space.

The population vectors 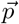 in phase space, whose components are the allele frequencies, lie only in the positive orthant of the phase-space and 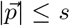. The hybridization process is assumed to take place suddenly in evolutionary terms so that the vector 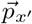 of an admixed population *x*′ is then a linear combination, in the vector algebra sense, of the contributor allele frequency vectors. For a 2-way admixture process we get 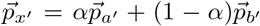, with *α* ∈ [0, 1] the contributing fraction of population *a*′ [10, 14]. However, no experimental information about primed populations is usually available and we have to pose the linear combination with the proxy populations

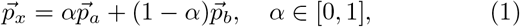

which is only exact at the time the admixed population is born.

Mathematically, the linear combination (1) stands for an over-determined linear algebraic system of dimension *s* with only one unknown, *α*. We can estimate *α* in the sense of Least Squares (LS) to give

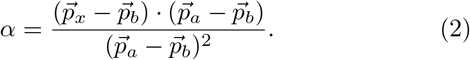

Here, the dot stands for the scalar product of two vectors

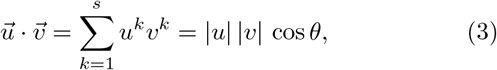

where *θ* is the angle between vectors 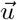 and 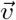, and *u*^*k*^ the *k*-component 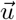. Alternatively, 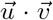 is the length of the projection of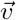along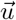, times 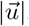.

For future reference it is convenient to remind the graphical interpretation of the LS solution (2) as the slope of the best fitting line through the origin to the set 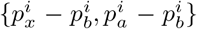 *i* = 1, …, *s*

It is straightforward to obtain, from (1), the constraint

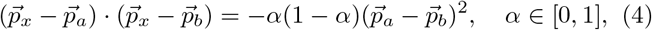

which conveys that the lhs must be negative and therefore the angle *ϕ* between vectors 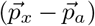 and 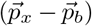, obtuse. This is the mathematical condition for admixture and in terms of scalar products reads

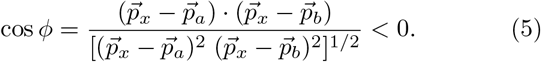

The angle *ϕ* at vertex *x* in the triangle Δ *axb* must be obtuse in a population admixture case.

The geometric interpretation of the *α* estimate (2) goes through the triangle Δ *axb* in Figure 1. The *α* value corresponds to the ratio whose numerator is the length of the orthogonal projection of the side 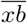 onto the side 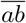, and the denominator is the side length 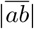. By now, this straightforward approach does not appear to benefit from the problem’s high dimensionality considerations mentioned above.

To pave the way for a connection with the *f*-statistics formalism, let us consider *k* auxiliary populations *i, j*, as those represented in the Figure 1. The difference vectors 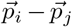 define arbitrary directions in phase space. Projecting the system (1) along them leads to the overdetermined system

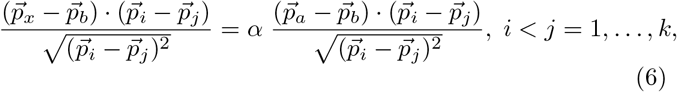

with *k*(*k* − 1)/2 equations and one unknown, *α*. The denominator ensures that the projection is properly done with unit vectors: 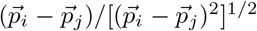. We can see (6) as the system (1) projected into a random subspace and, continuing the game, the LS solution reads

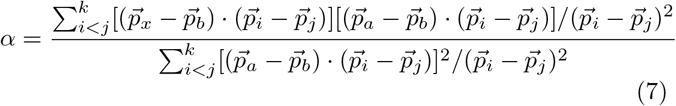

Indeed, the projection of the linear system (1) along the arbitrary directions 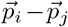 appears to be a whimsical procedure, raising the question: how much does the *α* estimate depend on the auxiliary populations choice? Or, what is the gain of using (7) instead of (2)? To address these and other issues some needed mathematical considerations [15] are explained next. The random projection (6) will prove to be an effective strategy and a basic element in the *f*-statistics approach to the population admixture problem.

### 2.1 Interlude: the blessing of dimensionality

The Johnson-Lindenstrauss (JL) lemma provides the first mathematical element we require. It states that a set of m points in a large dimensional space ℝ^*n*^ may be projected into a random subspace ℝ^*d*^, with *d* ≪ *n*, such that the pairwise distances of the point set are approximately preserved. The interesting point is that for a given pairwise distance accuracy the reduced dimension *d* is independent of the original dimension *n*. Only the number of points m has an impact.

#### Lemma 1

*Johnson-Lindenstrauss*. Given ϵ ∈ (0, 1/2), then for any set of points *S* = {*x*_1_, …, *x*_*m*_*}* in ℝ^*n*^, there exists the mapping *M* : ℝ^*n*^ *→* ℝ^*d*^ with *d* = *𝒪* (log(*m*)/ϵ^2^) such that

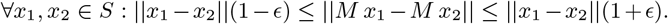

In algorithmic practice the mapping *M* is a *d × n* random matrix. We will associate the raws of this matrix to the difference vectors of auxiliary populations that carry out the projections in (6).

The second element refers to the quasi-orthogonality of random vectors in high dimension.

#### Lemma 2: quasi-orthogonality in high dimension spaces

Let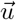 be a unit vector in *R*^*n*^, and 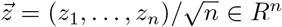, with each coordinate *z*_*i*_ at random from [−1, 1]. If *θ* is the angle between 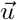 and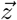, then we have the following bound on the probability

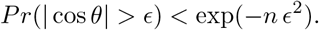

The amount of quasi-orthogonal random vectors grows exponentially with the space dimension. Figure 3 captures this phenomenon showing the histogram of the values of the cos *θ* between the vector (1, …, 1) and 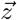 for four different values of the space dimension. The histogram shrinks as far as the dimension increases indicating quasi-orthogonality (cos *θ ≃* 0).

**Figure 3:**
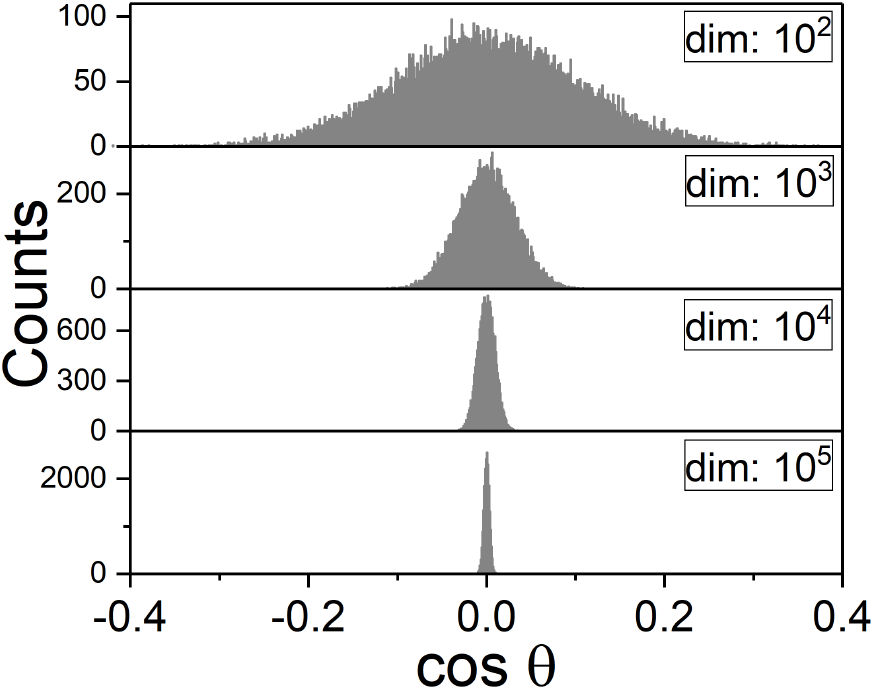
Histograms of 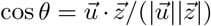, with 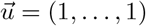, evaluated in four different dimensions. The distribution gets narrower as far as the space dimension increases.

In data mining, the JL lemma is used to reduce the numerical burden associated with large datasets. In our case the true utility stems from the involved projections and from the quasi-orthogonality property, which bring the proxies closer to the ancestral population admixture configuration in the allele frequency space by removing the post-admixture drift contribution. Given the populations *a, x, b* in Figure 1, we introduce the vector decomposition

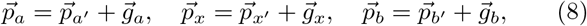

where 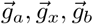, stand for the post-admixture drift contributions to the ancestral populations with each coordinate at random from [*−*δ, δ], and δ small. Next, we analyze the two scalar products in the numerators of (6) as regards quasi-orthogonality. For one of them

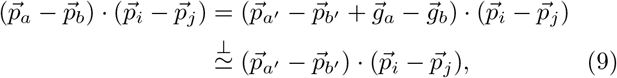

because lemma 2. And similarly for the other numerator. As a result, everything happens (approximately) in (6) as if we were working with ancestral populations rather than proxies. This is a feature already pointed out in the *f*-formalism using statistical foundations [1].

With *i, j* fixed (6) reduces to a scalar equation, which amounts a drastic dimensional reduction of the system (1) from dimension s ≫ 1 to a subspace of dimension d = 1, defined by the vector 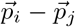. In the light of JL lemma, d = 1 must convey a significant distortion. By letting *i, j* be *k* > 2 running auxiliary populations, we define a higher dimension subspace of dimension *d* = *k*(*k*− 1)/2. The auxiliary pairs then define arbitrary directions to form the vector basis of a subspace, say the JL subspace. We expect less distortion after projection with *k* large enough (i.e., d > 1) than with *k* = 2 (i.e., *d* = 1). It is expected the approximate preservation of pairwise distances between vertices, combined with the approximate suppression of post-admixture drift contributions, to transform the phase space triangle Δ*abc* into a more obtusangle triangle 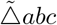 in the JL subspace, namely, a configuration close to an admixture at birth. It turns out that the very rule (2), when applied in the JL subspace, becomes (7).

### 2.2 The *f* -formalism: definitions

In terms of scalar products, the three basic *f*-statistics read

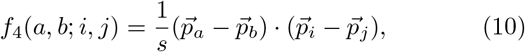

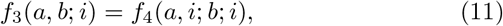

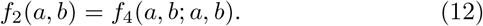

The three statistics are scalar products normalized to s, by definition. Note that f_2_ and f_3_ are degenerate cases of f_4_.

The first, *f*_4_(*a, b; i, j*), has a phylogenetic meaning. In the phylogenetic tree (((*i, a*), *b*), *j*), it determines the amount of shared drift between clades (a, b) and (i, j), termed *left* and *right* populations respectively. For *i, j* fixed, (6) reduces to a scalar equation for *α* that can be written down as

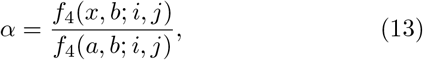

provided the denominator does not vanish. This estimate is the so-called *f*_4_ ratio and was used in the early stage applications of the *f*-statistics formalism as the basis to determine the 2-way admixture coefficient. In Section 2.6 the behavior of this estimate is worked out.

The treeness mathematical condition for the four populations phylogenetic tree above is

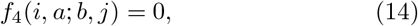

a constraint that will be used in the 3-way admixture case.

The second, *f*_3_(*a, b; i*), stands for the genetic distance between population *i* and the node (*a, b*), measured on the phylogenetic tree ((*a, b*), *i*).

The third, *f*_2_(*a, b*), is the squared Euclidean distance between a and b in phase space, normalized to s.

The estimate (2) can be written down in terms of *f*-statistics

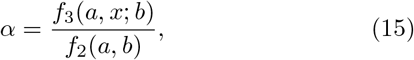

whereas (7) is given in the next Section. The constraint (5) is equivalent to f_3_(a, b; x) < 0, and reads

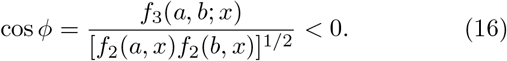

The f_2_ statistics is closely related to quasi-orthogonality in high dimension. Three populations that have diverged merely by genetic drift from a common root verify quasi-orthogonality in phase space: 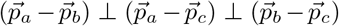, and for some adequate ordering the Pythagorean theorem in approximate form reads

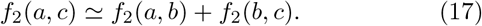

This identity is usually referred to as additivity of the f_2_ distance. Phylogenetic trees of populations that have evolved under pure genetic drift enjoy this property. Population admixture [16] as well as any event such as a bottleneck break additivity because the quasi-orthogonality requirements are not fulfilled.

### 2.3 Two-way admixture: the JL way

The system (6) is the JL dimensional reduction proposed for (1) in Section 2.1 and may be written down in terms of *renormalized f*-statistics

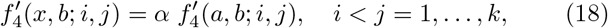

where we have defined

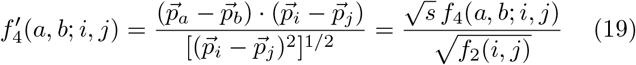

We refer to 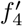 as the renormalized *f*_4_ because the *standard f*_4_ is a scalar product already normalized to the number s of SNP, by definition. Whereas in *f*_4_ the normalization coefficient is a constant, in 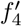 the normalization changes with the auxiliary pair. Whenever *k* > 2, the denominator of 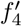 weights differently every equation in the system (18). Thus, distal auxiliary population pairs in phase space render less relevant that equation in the system with respect to the LS solution.

In agreement with the phylogenetic interpretation of *f*_4_, the system of d equations (18) can be interpreted as a *weighted proportionality law of the shared drift*. If the law is correct, a plot of the experimental values 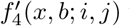 vs. 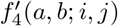 for several auxiliary populations yields points close to a straight line with slope *α*. The LS fit leads then to the remarkable formula

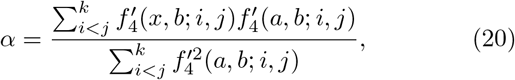

which collapses to the *f*_4_-ratio estimate (13) if we restrict ourselves to just one pair of auxiliary populations.

### 2.4 One triangle to rule them all

Given that the set 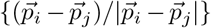 is a quasi-orthonormal basis, the sums in (20) represent scalar products in the JL subspace, in good approximation. This is because 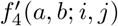 is the JL subspace component of 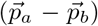 along the direction given by 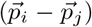. Therefore, the *α* estimates provided by (2) and (20) do share the same geometric interpretation. As explained above, given the triangle Δ *axb* in the allele frequency space, (2) is the ratio of the length of the projection of 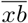 onto 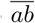, over 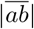. Concomitantly, (20) is the analogous ratio computed with the projected triangle, say 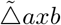, in the JL subspace. The advantage of (20) is that, according to the discussion in Section 2.1, the vertices of the JL projected triangle 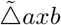 are expected to be almost a linear alignment, the characteristic of an admixture at birth. Moreover, we can extend this reasoning to estimate the angle φ at the vertex *x* of the JL projected triangle 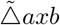:

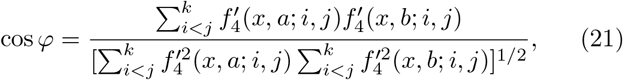

to be compared with cos *ϕ* in (16). Here, use has been made of the identity (3). As far as cos *φ* ≪ cos *ϕ*, or more precisely, as far as *φ*→ 180°, it is expected (20) to improve the *α* estimate with respect to the determination (2) or (15) in the full allele frequency space. We will show the substantial validity of these arguments in Section 3.

In the next section we connect the present derivation with that from standard *f*-statistics.

### 2.5 Two-way admixture: the *f* -statistics way

For the case of 2-way admixture, the current determination of *α* in the standard *f*-statistics framework is as follows. Let us rewrite the linear combination (1) as

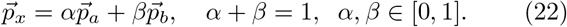

Then, due to the linear character of the scalar product we get

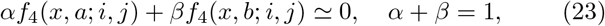

for any pair of auxiliary populations *i, j*. Considering d = *k*(*k* −1)/2 such pairs we get an overdetermined linear system with two unknowns and one constraint that can be formally written down as

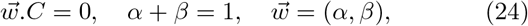

where C*∈* ℝ^2*×d*^. A solution of LS type could be worked out at this point. Instead, the matrix C is decomposed via Singular Value Decomposition as: C = A.B. This is the seed for an iterative process that minimizes the log-likelihood for (A, B) and takes into account the error introduced by the SNP sampling via a block jackknife resampling procedure. It is clear that no analytic expression can be given beyond this point. The admixture proportions are then found as the LS solution of

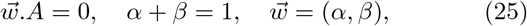

here A*∈* ℝ^2*×d*^. The special purpose *AdmixTools* package, and particularly the *qpAdm* function, is used for this task [17, 1, 6].

In order to obtain results that are comparable to those in Section 2.3, let us return to (23), that can be rewritten as

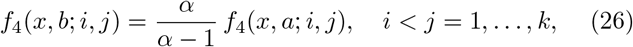

once the constraint has been explicitly introduced into the equations. A little algebra yields the system

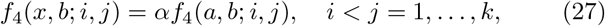

to be compared with (18). Namely, from the phylogenetic interpretation of *f*_4_ as shared drift, given the tree (((*i, a*), *b*), *j*) and the hybrid *x* with donors *a* and *b*, the shared drift between (*x, b*) and (*i, j*) must be proportional to the one between (*a, b*) and (*i, j*), where is *α* the proportion of population a contributing the admixture. The problem is then similar to that in (18). In a 2D representation, the set of values {*f*_4_(*a, b; i, j*), *f*_4_(*x, b; i, j*)}, with *i* < *j* = 1, …, *k*, should appear close to a straight line with slope α, as far as the admixture model is correct. The explicit LS formula to estimate the slope reads

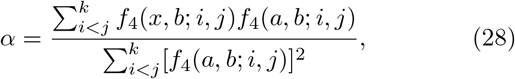

to be compared with (20). The *f*_4_ ratio definition (13) is recovered for *i, j* fix.

The LS solutions (24) and (25) are equivalent if the impact of the SNP sampling is not taken into account. In that context we can compare the two different rationales that have lead to (20) and (28). Both, the geometry-based scheme and the standard *f*-formalism, assume the same admixture proportion for every single SNP, i.e. equation (1). The JL projection leads to the equation system (18), which is solved by LS. The standard *f*-formalism derivation assumes, instead, the proportionality in the shared drift between the donors and the auxiliary pair, i.e. equation (27) which derives from (24).

Alternatively, we could interpret (27) as a JL-like projection which does not use unit base vectors. Therefore, unless all vectors 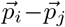 have similar modulus, the projection will suffer from a certain degree of distortion. A formal justification that alleviates this issue relies on a high-dimensional effect known as *concentration of the norm* [18]. It establishes that the norm of random vectors generated from the same distribution tends to a fixed value. In other words, the vectors cover approximately a hyper-sphere. Whether or not the vectors 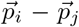 fit strictly the mathematical requirements, the outcomes of (18) and (27) might make little difference as regards the *α* determination. The numerical simulations buttress this argument.

Eventually, we have looked at the impact of SNP sampling using block jackknife resampling for a number of 2D realizations (27). As regards the determination of the mixture coefficient, these error estimates and those from the linear fits are of the same order in numerical simulations.

In the next section we interpret possible outcomes of the *f*_4_ ratio evaluations in the light of the JL dimensional reduction.

### 2.6 The *f*_4_ ratio illustrated

The evaluation of the *f*_4_ ratio (13) on its own corresponds to a projection of two sides of the triangle Δ *axb* in Figure 1 onto a one dimensional subspace defined by the difference vector of the two auxiliary populations. The geometry of such an evaluation is sketched in Figure 5 where, besides the triangle, four different one dimensional projections have been depicted. The diagram is planar only for the sake of clarity. The dashed segments 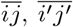 and 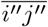 stand for directions defined by the auxiliary pairs 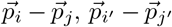 and 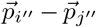. Gray lines are eye guides for the following orthogonal projections:

1. Projection on 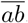. The sides 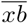 and 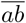 yield projection segments of lengths *p* (blue) and *q* (red), respectively; and *α* = *p/q* ∈ [0, 1], always in range under mathematical admixture conditions. This corresponds to case of (2), and is not *f*_4_ ratio.
2. Projection on 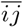. The sides 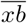and 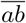 yield projection segments of lengths *s* (blue) and *r* (red), respectively. The corresponding *f*_4_ ratio yields the estimate *α* = *r/s* > 1, out of range.
3. Projection on 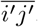. The sides 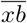and 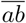 yield projection segments of lengths *t* (red) and *u* (blue), respectively; and *α* = t/u *∈* [0, 1], in range.
4. Projection on 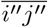. The sides 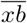 and 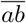 yield projection segments of lengths *y* (red) and *z* (blue), respectively; and *α* = − *y/z* < 0, out of range.

Projection#1 does not belong to the *f*_4_ ratio realm. It is given for the sake of completeness. It always leads to *α ∈* [0, 1], provided the admixture condition holds, namely *f*_3_(*a, b; x*) < 0. In cases with small post-admixture drift, which conveys an angle close to 180° at vertex *x*, equation (2) should provide an accurate determination. This scenario takes place in recent population mixtures.

Projections #2, #3 and #4 are characteristic of the *f*_4_ ratio procedure. Projection#3 gives a mathematically admissible estimate, whereas projection#2 and projection#4 fail to be in range.

In view of the JL lemma, it is understandable that the drastic dimensional reduction to dimension one has caused a great distortion in the projection. In the numerical simulation section we illustrate further on this issue.

### 2.7 The choice of auxiliary populations

The interpretation of the summations in (20) and (21) as approximate scalar products relies on the quasi-orthogonal character of the set of normalized difference vectors of auxiliary populations. Ideally, the auxiliary populations should provide an orthonormal basis but, in practice, it is only an approximation. This aspect will be illustrated in Section 3.3. Concomitantly, this vector basis should be as less orthogonal as possible to the vector 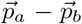, for a very important reason. In a 2D plot whose slope determines α, every point corresponds to an auxiliary pair *i*, j. Some pairs are irrelevant as regards the admixture problem, which happens whenever 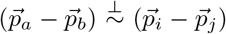, namely, small shared drift *f*_4_(*a, b; i, j*). For fixed donors, such an auxiliary pair does not contribute significantly to the *α* estimate because that point in the 2D plot remains close to the origin. Indeed, a straight line fit through the origin can be interpreted as a weighted average of the slopes defined by every single point, where the weight is the horizontal squared distance of that point to the origin^1^.

Therefore, auxiliary population pairs with large shared drift are the relevant ones.

The desideratum for a perfect auxiliary population pairs set is then: inner orthogonality and significant shared drift with donors (i.e., non orthogonality!). The practice goes otherwise. First, it is expected that some shared drift will be found among some of the possible combinations in the group of auxiliary pairs, leading to the quasi-orthogonality of the set. Second, configurations without shared drift with donors will unavoidably result from an unsupervised procedure to create pairs *i, j*. Fortunately, the corresponding points in the 2D plot will cluster quite close to the origin and thus should not significantly impair the fit’s slope accuracy.

### 2.8 Difficulties with introgression

When the contribution at admixture birth of (say) donor *b*′ is tiny, i.e. *α ≃* 1, complex situations are likely to arise in which genetic drift leads to misleading population configurations in phase space. This introgression situation is depicted in both panels of Figure 4 where the birth of the population admixture occurs along the segment 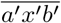 and it is meant that the contribution of *b*′ is tiny. The genetic drift is represented by the Brownian-like trajectories that drive *a*′, *x*′, *b*′ to the triangle Δaxb in phase space.

**Figure 4:**
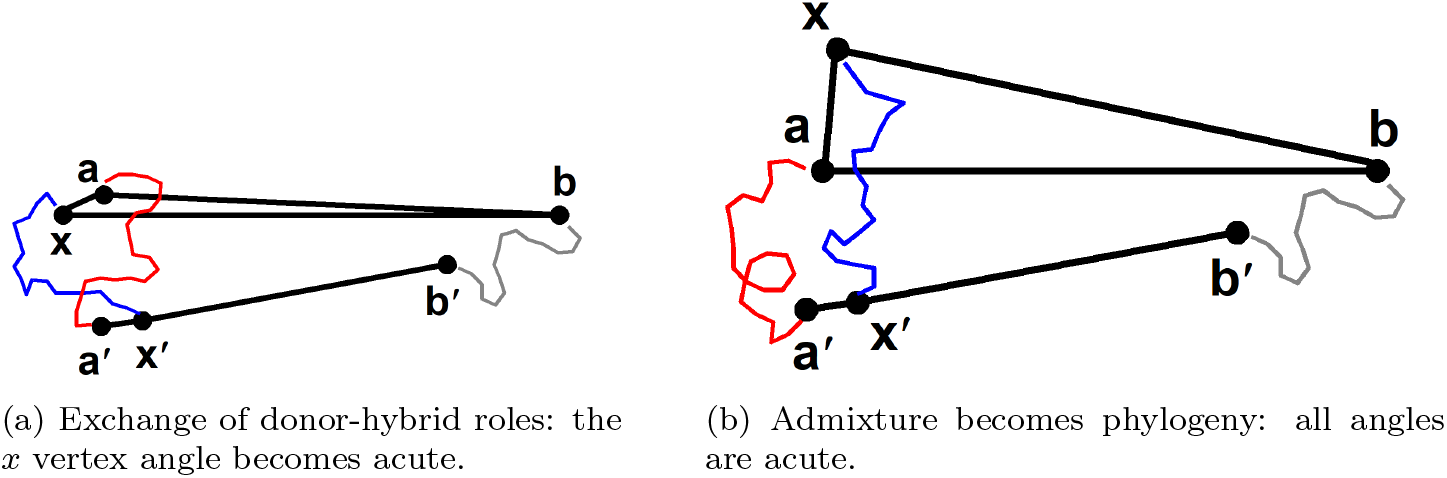
Planar representation of two introgression instances where genetic drift leads to misleading configurations in allele frequency space. Triangle Δ*axb* and segment 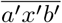 are as in Figure 1.

**Figure 5:**
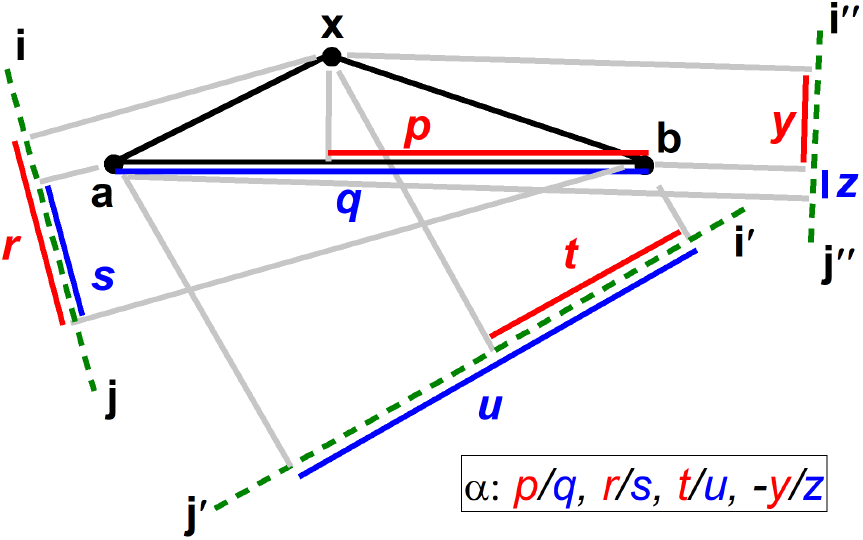
Planar representation for different estimates of the *f*_4_ ratio. Symbols p, q, r, s, t, u, y, z, stand for segment lengths. The ratio p/q is from (2), and r/s, t/u, *−*y/z are the *f*_4_ ratio (13) evaluated with three different pairs of auxiliary populations. Triangle Δ *axb* is from Figure 1. Dashed green lines are directions defined by auxi<u>lia</u>y pa<u>irs</u>. Gray lines are eye guides for orthogonal projections. Red and blue segments are projections of the triangle sides 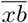 and 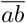, respectively.

In Figure 4a the initial hybrid *x*′ and the initial donor *a*′ are driven by genetic drift to the wrong vertex in the triangle Δ *axb*, in the sense that a appears as a hybrid and *x* as a donor, according to the admixture geometric condition.

In Figure 4b the Brownian-like trajectories are such that all the angles in triangle Δ *axb* are acute, indicating phylogenetic treeness rather than population admixture.

Both scenarios are likely to take place in long enough evolutionary histories. However, the closeness between *a*′ and *x*′ shorten the time lapse to occur. This qualitative observation can be bounded. The shortest trajectories that lead to the scenarios of Figure 4 are when *a*′ and *x*′ meet at midway of the segment 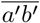 in phase space. The squared distance covered by both *a*′ and *x*′ is 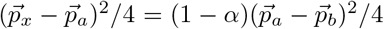. We can compare two population admixtures with the same three populations and different proportions, *α*_0_ and *α*, on the basis that the average squared distance covered in a Brownian motion is proportional to the time elapsed. Let t_0_ and t be those time intervals. A lower bound for the relative time in which the admixture test is not fooled reads then

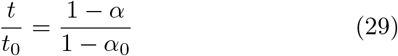

If we compare introgression (1 − *α* ≪ 1) with respect to a balanced admixture (say, *α*_0_ ≃ 0.5), it turns out that *t* ≪ *t*_0_: the time interval to fool the admixture test is much shorter in introgression than in a balanced admixture.

To what extent, in an introgression scenario, the JL projection will yield a proper admixture triangle 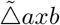 in the subspace is an open question. Some numerical experiments are presented bellow.

### 2.9 Three-way admixture

The JL projection procedure may be generalized to a population admixture with more than two donors. Next we illustrate the 3-way case with contributors a, b, c. In the case when the three parental are related by phylogenetic treeness, a necessary mathematical condition for *x* to be a hybrid is that the four populations *{*a, b, c, x*}* be not a phylogenetical tree, namely

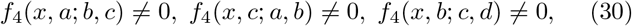

where we have used (14). Under the hypothesis that all the s allele frequencies are combined with fixed proportions, the linear combination now reads

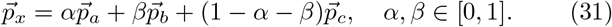

The four points involved form a planar configuration in phase space at admixture birth with the hybrid inside a triangle of contributors (see 4). The effect of the post-admixture drift makes the co-planar structure in phase space to become an irregular tetrahedron.

After JL projection with the *d* = *k*(*k* − 1)/2 pairs of auxiliary populations the system (31) reads

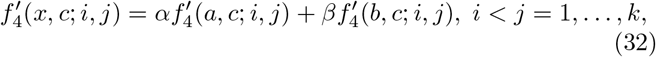

Generalizing the result in Section 2.3, a way to look at the LS solution of the system consists in finding the best fit in 3D to a plane through the origin with the d points in the set 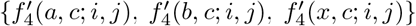. Explicit formulas for *α* and β are

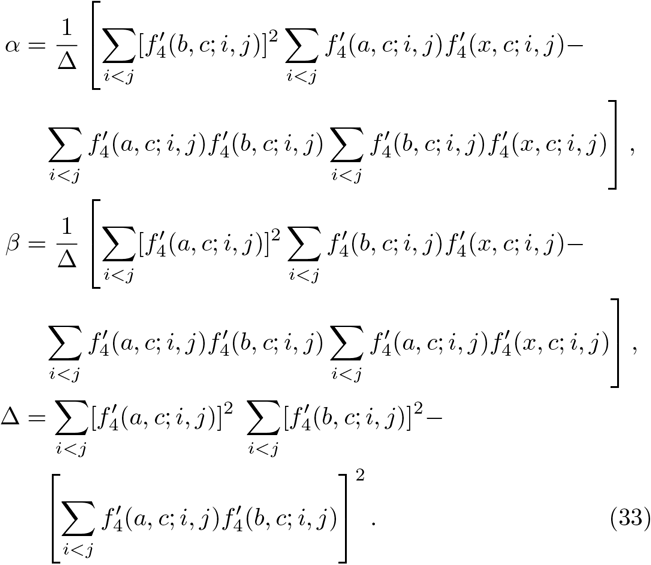

The alignment of the three populations at a 2-way admixture birth has the 3-way admixture analogous in the co-planar character of the four populations involved, for which an explicit proof may be found in 4. It would be convenient to have an index to measure to what extent the post admixture tetrahedron gets flattened in the JL subspace and therefore similar to an admixture at birth. To this end, we have computed the tetrahedron volume V, the base area A and the height *h* (from base to hybrid vertex) in both the full phase space and the JL subspace. In order to have a dimensionless quantity we have defined a heuristic *flatness index* as the ratio of the tetrahedron height over the square root of its base area: 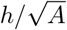. When the tetrahedron is not enough warped that the *x* vertex orthogonal projection falls outside the base, this indicator should be useful. Once more, the introgression regime is more sensitive to this circumstance.

The tetrahedron volume and the area have been calculated with the Cayley-Menger determinant and the Heron formula, respectively [18]; which use just the edge lengths, namely 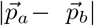, between two population vertices a and b. Explicit formulas are given in 4 where it is shown that the flatness index reads 3V/A^3*/*2^.

## 3 Numerical simulations

We have run a number of simulations using real-world data to illustrate the formal arguments presented above. To this end, we have used the population dataset *v37*.*2. 1240K HumanOrigins* from Reich Lab [19] to build up a dataset of forty populations: English, Yoruba, Biaka, Mbuti, French, Druze, Sardinian, Italian North, Basque, Adygei, Nganasan, Lithuanian, Lebanese, Han, Papuan, Surui, Karitiana, Aleut, Ju hoan North, Gujarati, Iranian, Australian, Khomani, Yukagir, Chukchi, Ulchi, Tubalar, Koryak, Even, Brahui, Balochi, Hazara, Makrani, Pathan, Kalash, Burusho, Japanese, Orcadian, Mayan and Russian. The effective number of SNP is s = 597.568.

We have built up hybrid populations as in the linear combinations (1) and (31) taking two, or three, populations from the set whereas the remaining populations in the dataset serve as auxiliary populations. To simulate post-admixture drift, noise of amplitude ∈ ϵ [0, 0.5] has been added to the allele frequencies of the donors and the hybrid. The quantity added is ϵδ, with δ ∈ [−1, 1] a random number.

### 3.1 Results for 2-way admixture

We commence with the simulation: 0.1 English + 0.9 Yoruba, and large noise amplitude ϵ = 0.5; namely a long time-evolution after hybrid birth, in order to enhance the misalignment of points in 2D plots.

Figures 6a and 6b show the outcomes of the linear relationship suggested by (18) and (27), respectively. The solid line has slope 0.1, the nominal admixture proportion whereas the fits give estimates *α* = 0.105 and 0.106 for the 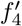 and *f*_4_ representations. Henceforth, the admixture proportions are given with the number of significant digits. The angle at the hybrid vertex in phase space is *ϕ* = 67.9°, which breaks the admixture condition. Even though, in the JL subspace the projected angle reads φ = 175.8°, which witnesses a good reconstruction of the geometric configuration at the admixture birth because it is close to 180°. The inset allows to appreciate, for points with small shared drift, that the slopes defined by every single point yield a large range of *f*_4_ ratio evaluations, most of then out of range.

**Figure 6:**
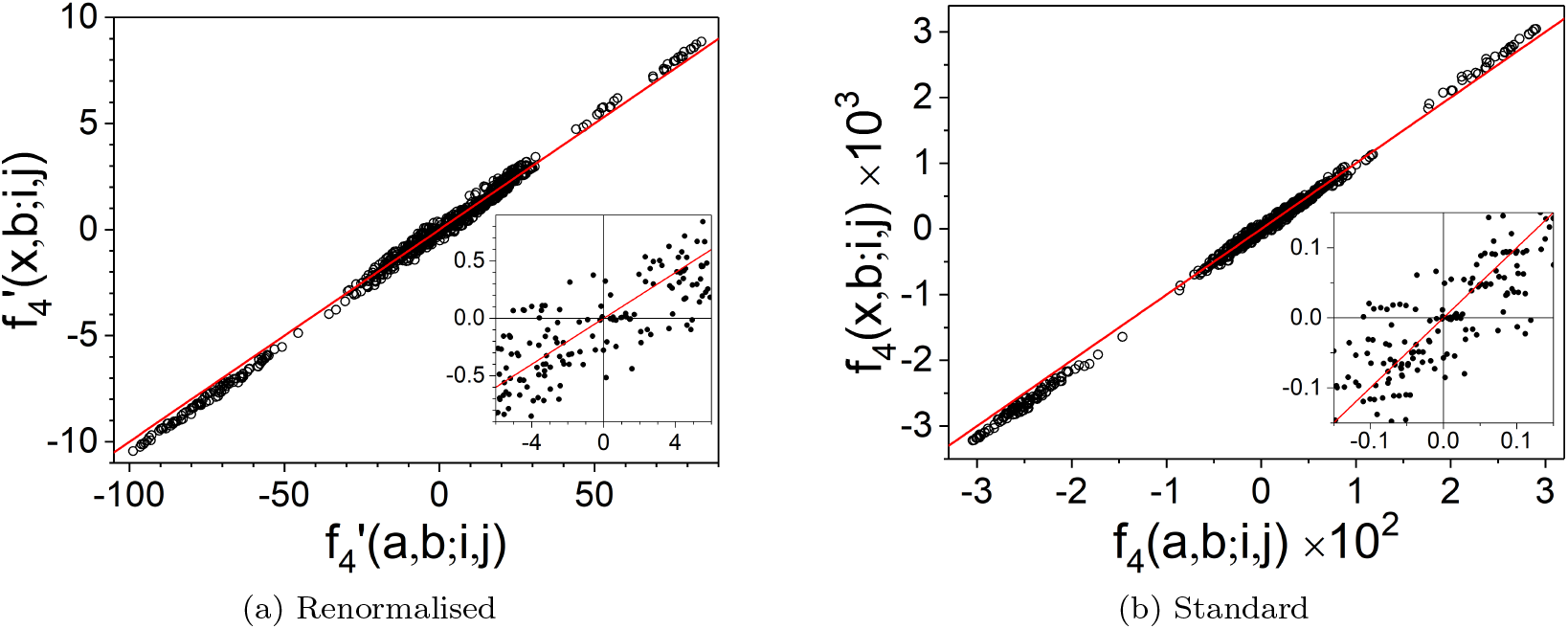
2D representations of population admixture. Simulated model: 0.1 *·*English + 0.9 *·*Yoruba, with noise amplitude ϵ = 0.5. Panel (a) is for the renormalized representation (18) and panel (b) is for the standard representation (27). Solid lines have slope 0.1. The zoomed area allows to appreciate the diversity of slopes one can get in an *f*_4_ ratio evaluation where the denominator is small by considering the slope defined by every single point.

Figures 7a and 7b are sanity checks and show real data outcomes for unlikely population admixture models: Sardinian = *α ·*Italian_ North +(1 *−* α) *·*Basque, and Italian North = *α* Basque +(1 α) *·*Adygei, respectively. The dispersion of the points is high in both panels. In the model of Figure 7a the estimate reads *α* = 0.98, in range. However, the angle *ϕ* = 59.1° fools the admixture test and points strongly out towards an unlikely admixture. The confirmation comes from the projected angle whose value is *φ* = 47.1° ≪ 180°, which is even worse than ϕ. In the model of Figure 7b the estimate is *α* = 0.80. The angle *ϕ* = 69.4° breaks the admixture test too. Although the projected angle *φ* = 115.5° does not, it is far from the required 180° in a proper reconstruction.

**Figure 7:**
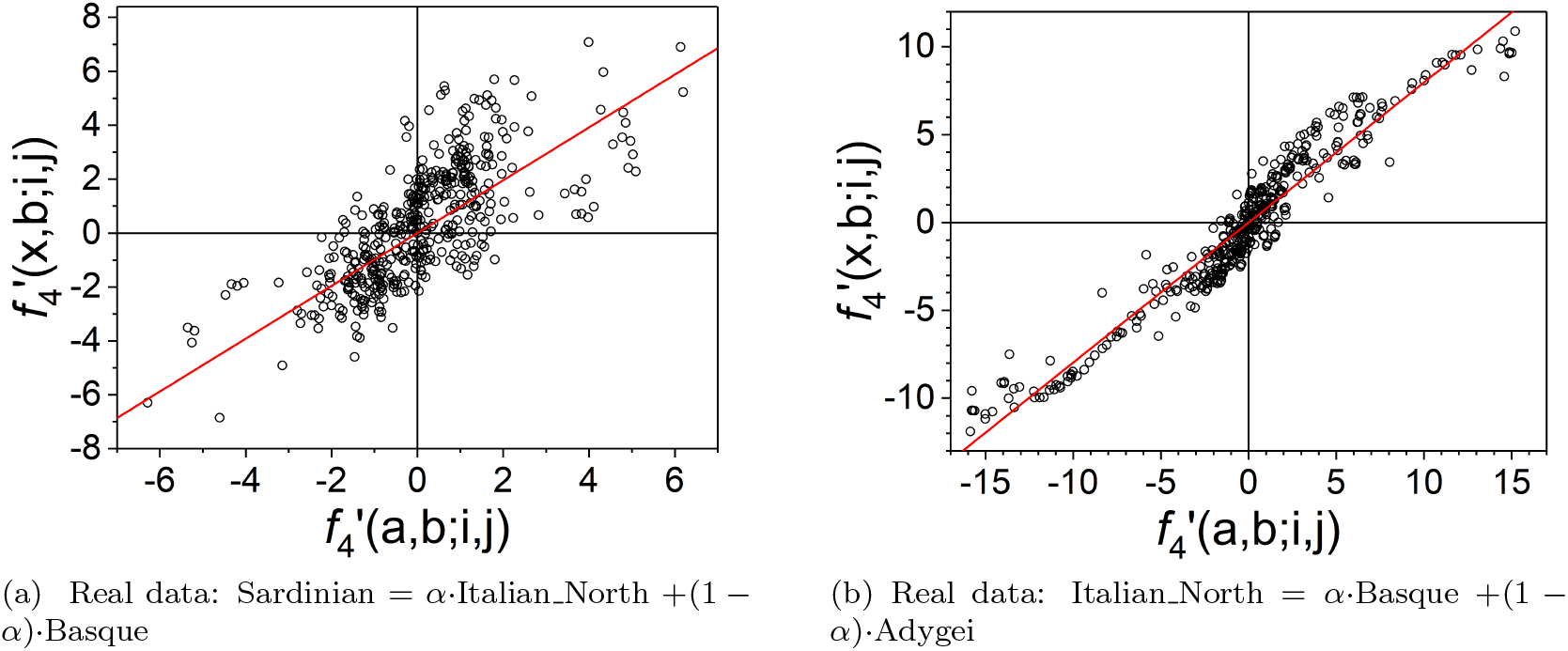
2D representations of population admixture. Two different unlikely real data models. Solid lines are linear fits through the origin with slopes (a) 0.98 and (b) 0.80.

The results of calculating the *f*_4_ ratio for each auxiliary pair are shown in Figure 8. The shared drift in absolute value is compared to the *f*_4_ ratio, i.e. the denominator as a function of the ratio. Figure 8a illustrates, with the simulated model: 0.1*·*English+0.9*·*Yoruba, that little shared drift leads commonly to unreliable *f*_4_ ratio estimates, a fact that the linear regression avoids. Concomitantly, for the model: Italian North = α*·*Basque +(1 *−* α)*·*Adygei, Figure 8b shows that unrealistic population admixtures lead to *f*_4_ ratio estimates of broad spectrum. The zoomed area in the inset has a similar scale to Figure 8a, for the sake of comparison. Both plots point out a concentration of the *f*_4_ ratio for shared drift large enough albeit the dispersion is much greater for the unrealistic admixture.

**Figure 8:**
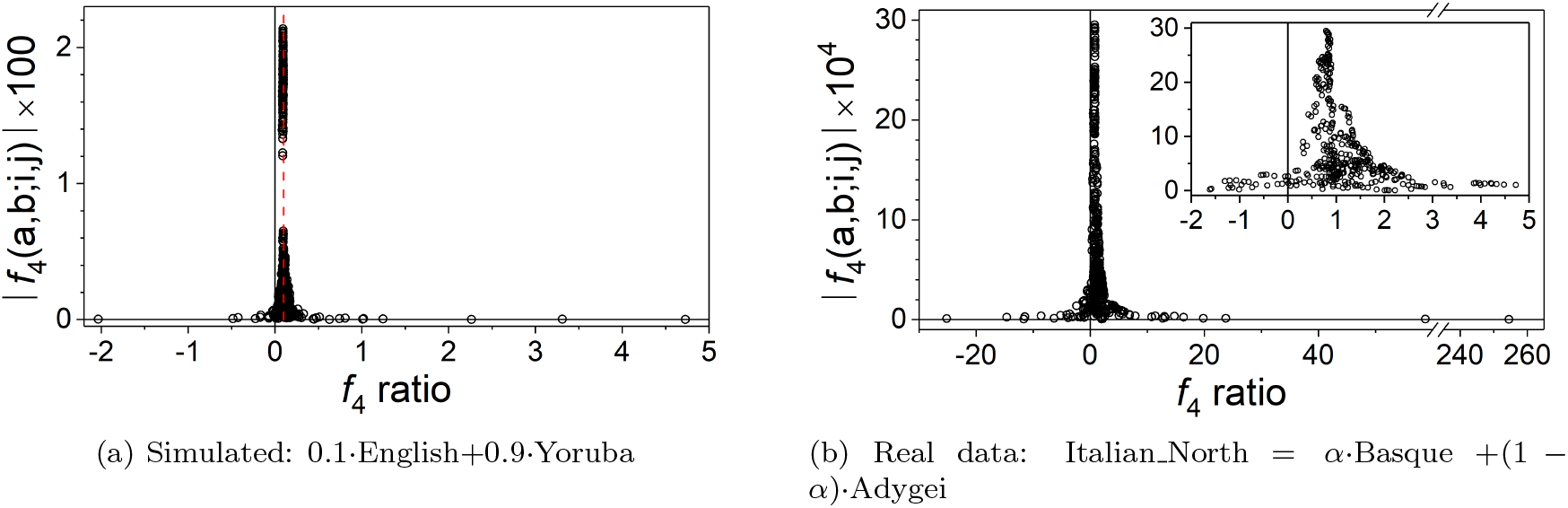
Behaviour of the *f*_4_ ratio values with respect to the shared drift. Panel (a) is for the simulated case 0.1·English+0.9 *·* Yoruba, and ϵ = 0.5. The dashed line stands for the nominal *α* = 0.1. Panel (b) is for the unlikely real data model: Italian _North = *α ·*Basque +(1 *−* α) *·*Adygei. The zoomed area in the inset has the same horizontal scale as panel (a).

Numerical outputs for simulations with various *α* values and noise amplitudes can be found in Table 1. Both, renormalized and standard approaches agree within three significant digits in the *α* estimate in almost all cases. The results find that irrespective of the noise amplitude (i.e., time elapsed since admixture) both the standard and the renormalized *α* estimates are robust Columns with the values for cos *ϕ* and *ϕ* itself are given. Notice that some cases fool the admixture test. Interestingly, the method still provides good estimates in such cases. As evidence of how near to the ancestral alignment the JL projection left the allele frequency vectors, the values of the angle φ approach 180°, witnessing a proper admixture reconstruction. In this regard, there is a tendency for φ to worsen as the noise amplitude increases: the ancestral admixture reconstruction is harder for long time-evolution.

**Table 1:** Simulations for the model: *α* English + (1 *−* α) *·* Yoruba, with various nominal *α* values and noise amplitudes ϵ. Columns for the renormalized (20) and the standard (28) approaches are given. The angles *ϕ* and φ refer to equations (16) and (21), respectively. Boldfaced data stand for cases fooling the mathematical admixture test (16). *Proxy* and *Ancestral* refer to preand post-JL projection, respectively.

| Nominal<br>$\alpha$ | Renorm.<br>$\alpha$ | Standard<br>$\alpha$ | Noise<br>$\epsilon$ | Proxy<br>$\cos \phi$ | Proxy<br>$\phi$ (deg) | Ancestral<br>$\varphi$ (deg) |
| --- | --- | --- | --- | --- | --- | --- |
| 0.1 | 0.099 | 0.099 | 0.1 | -0.55 | 123.7 | 179.8 |
| 0.3 | 0.299 | 0.300 |  | -0.87 | 150.7 | 179.9 |
| 0.5 | 0.503 | 0.503 |  | -0.91 | 155.0 | 179.9 |
| 0.1 | 0.100 | 0.100 | 0.3 | **0.13** | **82.6** | 178.9 |
| 0.3 | 0.297 | 0.297 |  | -0.06 | 93.5 | 179.6 |
| 0.5 | 0.497 | 0.497 |  | -0.13 | 97.4 | 179.7 |
| 0.1 | 0.105 | 0.106 | 0.5 | **0.38** | **67.9** | 175.8 |
| 0.3 | 0.305 | 0.305 |  | **0.34** | **70.3** | 178.1 |
| 0.5 | 0.497 | 0.497 |  | **0.32** | **71.2** | 178.4 |

Further admixture simulations, 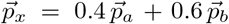, are shown in Figure 9 with the donors specified in the figure legend. The 2D fit estimates are very accurate. Notice that although the set of auxiliary population is very similar in all simulations, the plots exhibit a variety of scale ranges as a consequence of the diversity in the shared drift between the auxiliary pairs and the contributors.

**Figure 9:**
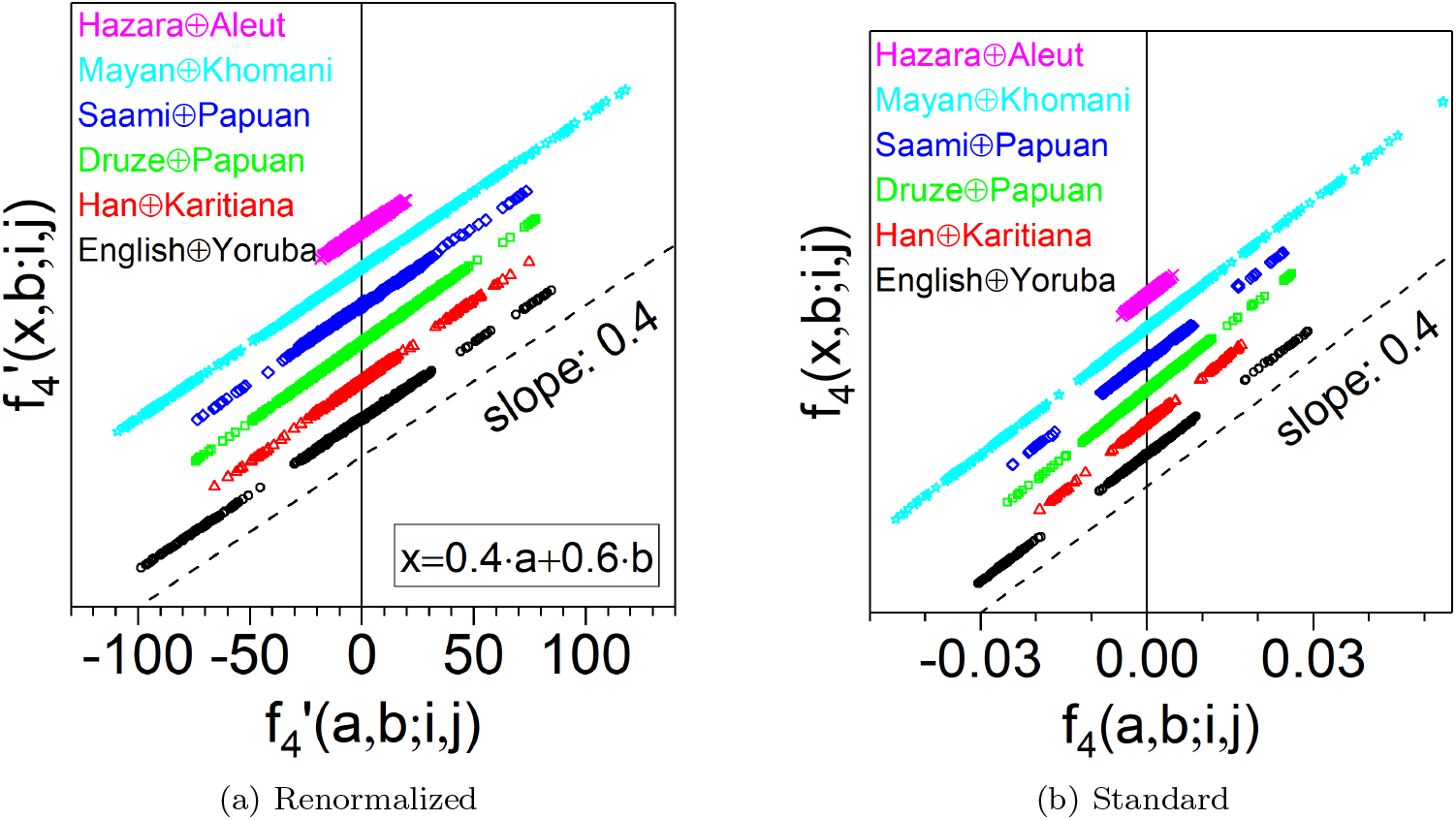
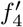 and *f*_4_ representations of the six different 2-way admixture simulations given in the legend, following the model *x* = 0.4a + 0.6b. Every data alignment in the plot corresponds to one hybrid, contains 703 points (auxiliary population pairs *i*, j) and has been vertically shifted for readability. The dashed line has slope: 0.4, and is given for visual reference. The legend is ordered as the alignments sequence.

We performed a jackknife study to see what impact sampling of auxiliary populations has on the determination of slopes. The results show that the effect is much less significant than the indeterminacy of the linear fits themselves.

### 3.2 Introgression

The drawbacks with introgression analyzed in Section 2.8 are brought to light in Table 2. We have simulated the admixture: *α*·English + (1 − α) *·*Yoruba, with small *α* and three different noise amplitudes. The agreement with the nominal values worsens with decreasing nominal *α* and increasing noise amplitude. For a given noise amplitude, the values of angles *ϕ* and φ find that a small *α* model is much more affected by post-admixture drift. Increasing the noise amplitude enhances this effect. The worst case represented is with *α* = 0.002 and ϵ = 0.3, namely, extreme introgression and long time evolution since admixture.

**Table 2:** Introgression regime for the model: *α ·* English + (1 *−* α) *·* Yoruba, i.e. nominal *α ≪* 1. Columns are coded as in Table 1, except boldfaced data which here stand for the cases compliant with the mathematical admixture test (16). *Proxy* and *Ancestral* refer to preand post-JL projection, respectively.

| Nominal<br>$\alpha$ | Renorm.<br>$\alpha$ | Standard<br>$\alpha$ | Noise<br>$\epsilon$ | Proxy<br>$\cos \phi$ | Proxy<br>$\phi$ (deg) | Ancestral<br>$\varphi$ (deg) |
| --- | --- | --- | --- | --- | --- | --- |
| 0.002 | 0.002 | 0.002 | 0.1 | 0.05 | 87.3 | 172.0 |
| 0.010 | 0.010 | 0.010 |  | **-0.01** | **90.8** | 178.4 |
| 0.050 | 0.049 | 0.049 |  | **-0.30** | **107.3** | 179.6 |
| 0.002 | 0.003 | 0.003 | 0.2 | 0.15 | 81.5 | 160.7 |
| 0.010 | 0.011 | 0.011 |  | 0.13 | 82.8 | 174.8 |
| 0.050 | 0.050 | 0.050 |  | 0.02 | 89.1 | 178.9 |
| 0.002 | 0.003 | 0.003 | 0.3 | 0.25 | 75.5 | 151.2 |
| 0.010 | 0.011 | 0.011 |  | 0.24 | 76.1 | 171.2 |
| 0.050 | 0.051 | 0.051 |  | 0.19 | 79.1 | 178.0 |

All this said, it is interesting to note that although the admixture criterion cos *ϕ* < 0 is only met in two out of nine simulations, the co-linear configuration at admixture birth in the JL subspace is nonetheless roughly obtained, because the estimations for the ancestral angle φ are manifestly obtuse in all cases.

### 3.3 The quasi-orthogonality of the dataset

The quasi-orthogonality of the set of auxiliary populations is a fundamental piece of the framework. Thus, the question of how much orthogonal the auxiliary population set is arises. The answer is in Figure 10 (bottom panel) which presents a histogram of the absolute value of the cosine of the angle between two population pairs. The distribution is peaked at 90°.

**Figure 10:**
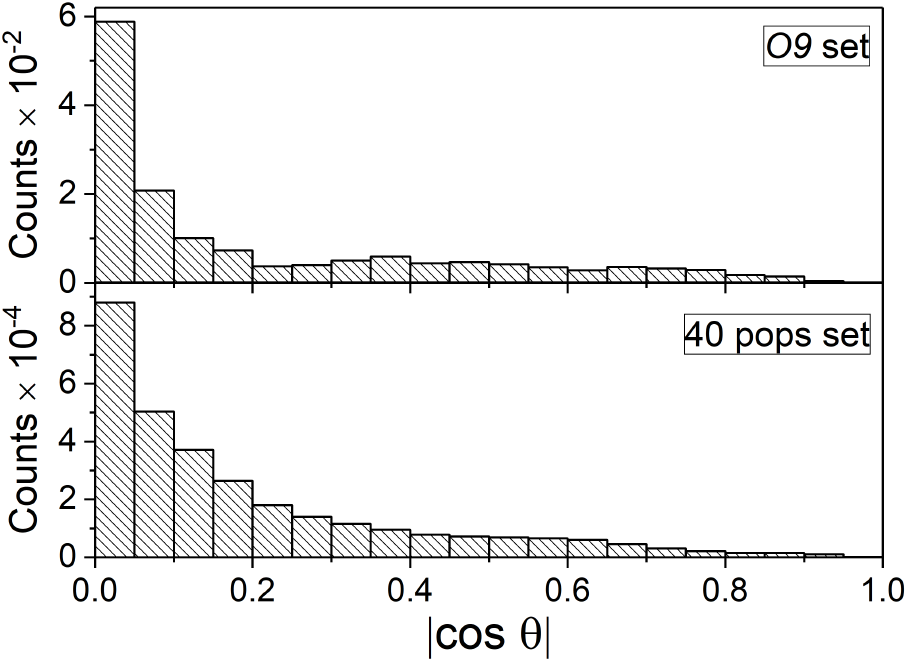
Quasi-orthogonality in allele frequency space. Histogram of the absolute values of the cosine of the angle *θ* between two auxiliary pairs. Top and bottom panels are, respectively, for the **O9** and the 40 populations dataset.

To buttress that this feature is not a coincidence we have generated a similar histogram associated to the so-called auxiliary population dataset **O9** [3]: Ust Ishim, Kostenki14, MA1, Han, Papuan, Onge, Chukchi, Karitiana and Mbuti. This gives 36 pairs of auxiliary populations and the effective number of SNP is in this case s = 380.574. The populations in this set have been carefully chosen in [3] according to their location in a phylogenetic tree, in contrast with the choice of populations in the 40 populations dataset which was rather whimsical. The quasi-orthogonality of the **O9** set is disclosed in the histogram of Figure 10 (top panel).

### 3.4 Standard versus renormalized *f*_4_ representations

The observed equivalence of the 2D representations, *f*_4_ and 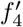, with respect to the *α* estimation points out that the implicit denominators in the equation system (18) cannot have a great relative diversity in magnitude. This is witnessed in the histogram of the Euclidean distance of the population pairs in the dataset given in Figure 11. The Euclidean distance of the bulk of pairs is in the range 144*±* 50, and the tails of the distribution are small. The variability of denominators is smaller than one order of magnitude.

**Figure 11:**
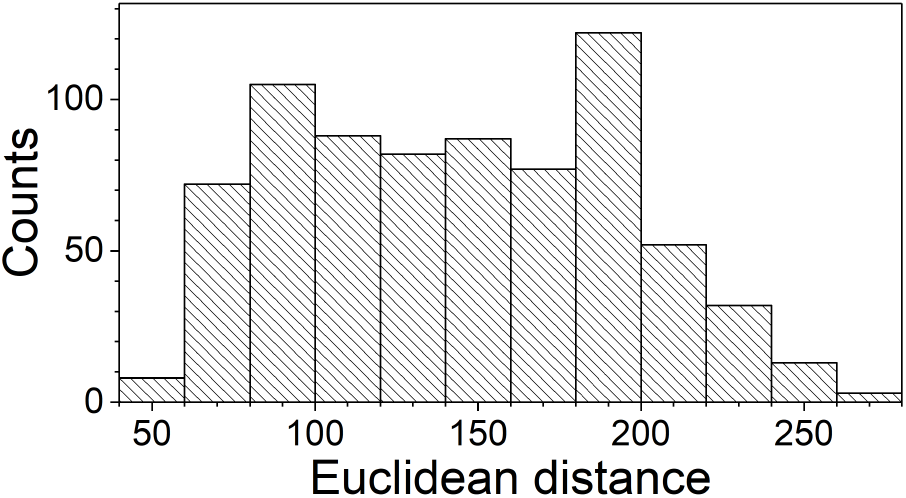
Histogram of Euclidean distances 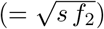 provided by auxiliary population pairs in the dataset.

### 3.5 Results for 3-way admixture

Results for the numerical simulation of 3-way admixtures are displayed in the four panels of Figure 12. The simulated model: 0.2*·* English +0.5*·* Yoruba +0.3*·* Biaka, is shown in panels (a) and (b) with two different perspectives. They exhibit the co-planar distribution of points expected from (31). The flatness index decreases from 0.501 to 0.017, i.e. one order of magnitude.

**Figure 12:**
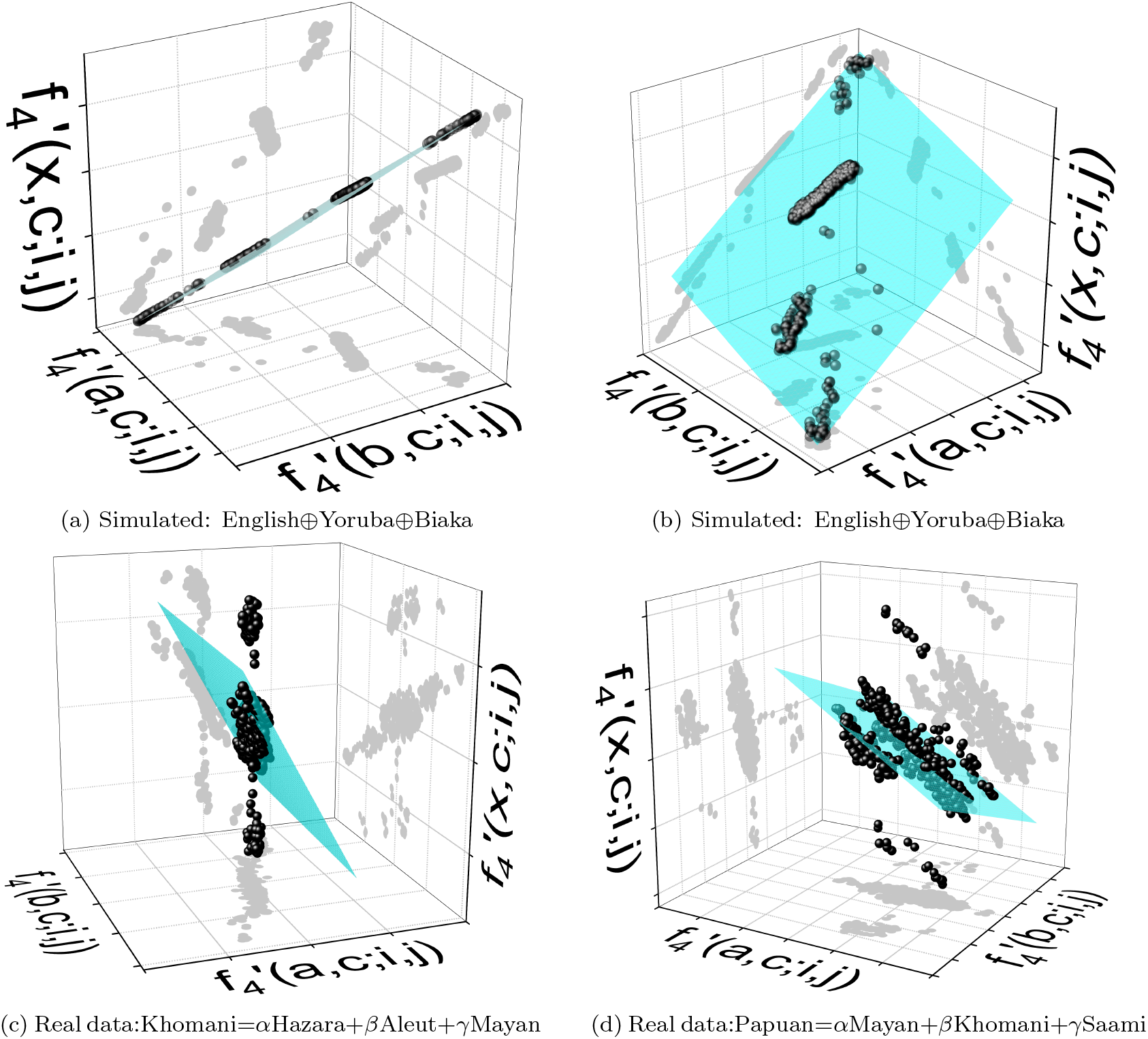
Three-way admixture. Panels (a) and (b) are for the admixture model generated by: *α* ·English+*β* · Yoruba+ *γ* · Biaka, with nominal *α* = 0.2, *β* = 0.5, *γ* = 0.3. Noise of amplitude *ϵ* = 0.1 has been added to the four populations prior to the fit. The fit gives admixture proportions: *α* = 0.18, *β* = 0.54, γ = 0.28. Panel (c) is the result of the unlikely mixture: Khomani = *α ·*Hazara +*β* · Aleut +*γ* ·Mayan. The fit outcome is: *α* = 1.6, β = *−* 0.1, γ = *−* 0.5. Panel (d) is the result of the unlikely mixture: Papuan = *α* Mayan +β Khomani +*γ* Saami. The fit outcome is: *α* = 0.46, *β* = 0.20, *γ* = 0.34. The planes are LS fits to data, and the gray dots are the orthogonal projections of data.

Table 3 lists results for simulations using different noise amplitudes and admixture proportion combinations. The first observation is that the determination of the coefficients is less accurate in the 3-way case than in the 2-way case (less significant digits). Thus the plane tilt in 3D is subject to more variability than the straight line slope in 2D. The cases likened to introgression, say α, β or γ = 0.1, tend to give the worst estimates. As a rule of thumb, the flatness index decreases between one and two orders of magnitude after the JL projection.

**Table 3:** Three-way estimates for fifteen simulations: *α ·* English + β *·* Yoruba + γ *·* Biaka. *Proxy* and *Ancestral* refer to preand post-JL projection, respectively.

| Noise<br>$\epsilon$ | Nominal | | | Linear fit | | | Flatness index | |
| --- | --- | --- | --- | --- | --- | --- | --- | --- |
| | $\alpha$ | $\beta$ | $\gamma$ | $\alpha$ | $\beta$ | $\gamma$ | Proxy | Ancestral |
| 0.01 | 0.2 | 0.5 | 0.3 | 0.191 | 0.505 | 0.305 | 0.209 | 0.008 |
| 0.10 |  |  |  | 0.182 | 0.500 | 0.319 | 0.501 | 0.017 |
| 0.30 |  |  |  | 0.151 | 0.640 | 0.209 | 0.952 | 0.022 |
| 0.01 | 0.1 | 0.7 | 0.2 | 0.094 | 0.705 | 0.201 | 0.268 | 0.073 |
| 0.10 |  |  |  | 0.088 | 0.698 | 0.214 | 0.524 | 0.012 |
| 0.30 |  |  |  | 0.072 | 0.814 | 0.114 | 0.966 | 0.023 |
| 0.01 | 0.8 | 0.1 | 0.1 | 0.797 | 0.107 | 0.096 | 0.421 | 0.009 |
| 0.10 |  |  |  | 0.797 | 0.102 | 0.101 | 0.583 | 0.013 |
| 0.30 |  |  |  | 0.739 | 0.226 | 0.035 | 0.981 | 0.026 |
| 0.01 | 0.5 | 0.4 | 0.1 | 0.497 | 0.404 | 0.099 | 0.324 | 0.008 |
| 0.10 |  |  |  | 0.497 | 0.397 | 0.106 | 0.524 | 0.013 |
| 0.30 |  |  |  | 0.457 | 0.523 | 0.020 | 0.949 | 0.020 |
| 0.01 | 0.1 | 0.1 | 0.8 | 0.075 | 0.112 | 0.813 | 0.389 | 0.010 |
| 0.10 |  |  |  | 0.054 | 0.105 | 0.841 | 0.602 | 0.024 |
| 0.30 |  |  |  | 0.007 | 0.306 | 0.686 | 0.991 | 0.033 |

As sanity checks, in Figure 12c we have tested the mixture: Khomani = *α* Hazara +*β* Aleut +*γ* Mayan, and in Figure 12d: Papuan = α*·* Mayan +β *·*Khomani +*γ* · Saami, with *α* + *β* + *γ* = 1. These farfetched population combinations display a distribution of points far from co-planarity. The fitted proportions are: *α* = 1.6, *β* = −0.1, *γ* = −0.5, for Figure 12c; and *α* = 0.46, *β* = 0.20, *γ* = 0.34, for Figure 12d. Interestingly, the flatness index in Figure 12c, where the admixture proportions are out of range, increases from 2.3 to 4.8 after JL projection. In Figure 12d, where the estimated proportions are in range, the index decreases moderately from 1.3 to 0.7. For the specific model: 0.2*·* English +0.5 *·*Yoruba +0.3 *·*Biaka, simulated with different noise amplitudes, the Figure 13 shows the volume and the flatness index of the tetrahedron in the full phase space and in the JL subspace. Every point stands for an average of 50 simulations. A substantial decrease in the tetrahedron volume is observed from full phase space (black squares) to JL subspace (red dots). This volume shrinking is not due to a mere scale change following the JL projection. Rather, the tetrahedron becomes flatter as a consequence of the very JL projection. This is proven by the flatness index curves which show the drastic flattening carried out by the JL projection albeit it modulates as far as the noise amplitude increases.

**Figure 13:**
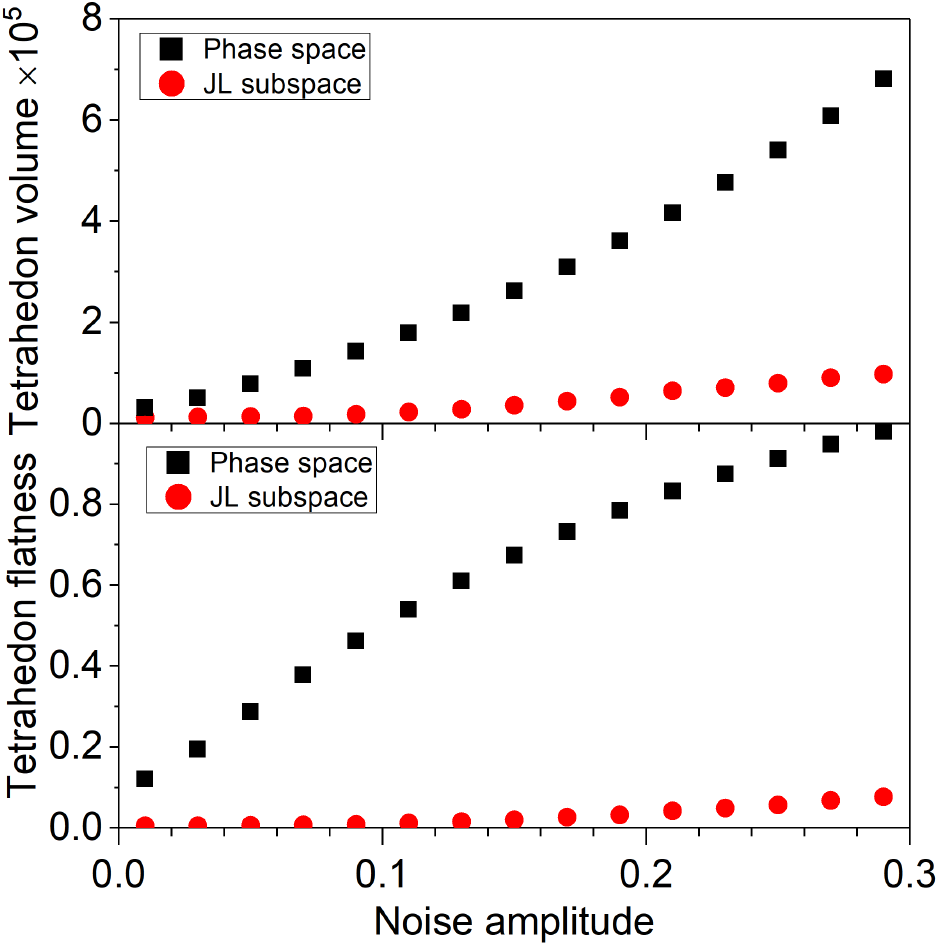
Three-way admixture: tetrahedron features as a function of the noise amplitude. Every dot stands for the average of 50 simulations with the model 0.2 *·* English+0.5 *·* Yoruba+0.3 *·* Biaka. One standard deviation error bars are smaller than the symbols. Upper panel: tetrahedron volume in the allele frequency phase space (squares) and in the JL subspace (circles). Bottom panel: flatness index in the allele frequency phase space (squares) and in the JL subspace (circles).

## 4 Discussion

Following is succintly described the essence of the geometric perspective about the *f*-formalism approach to the population admixture problem.

There is a formal part in which the populations in the dataset are viewed as points in the allele frequency space. The proxies of the admixture data configure a triangle or a tetrahedron in phase space, following the 2-or 3-way case. The auxiliary populations, in turn, define the random subspace where the proxies are projected and, according to the JL lemma, the relative proportions of the triangle or the tetrahedron are approximately preserved. The key point is that, due to the high dimension of the phase space and the random nature of genetic drift, the very JL projections deplete the post-admixture drift. Thus, once projected, the triangle and the tetrahedron become geometric configurations that are close to an admixture at birth.

In a 2-way admixture the proxies form a triangle in phase space and the mathematical condition for admixture reads 90° < *ϕ* < 180°, for the angle at the hybrid vertex. After JL projection, the new triangle should have the vertices almost aligned. The closer the new angle *φ* to 180°, the better the reconstruction of the native admixture configuration will have been. Moreover, numerical experiments have shown that cases with *ϕ* < 90°, and so outside the mathematical admixture domain, can still be successfully mapped to *φ* ∼ 180°. This is a new feature of the *f*-formalism that emerges in the geometric picture.

In a 3-way admixture, the proxy samples form an irregular tetrahedron in phase space which after JL projection becomes flatter. The closer to a co-planar configuration, the better the reconstruction will have been. The mapping efficiency is quantified by the heuristic flatness index.

The practical task reduces to carry out an appropriate linear fit. We are given a dataset with allele frequencies of a number *k* of populations, for a large number s of SNP. One population is the hybrid and two, or three, the contributors. The remaining *k −* 3, or *k −* 4, are used as auxiliary populaions. The admixture proportion estimates come from a linear fit through the origin to the set 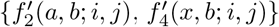 with (k*−*3)(k*−*4)/2 points, in a 2-way admixture. In a 3-way admixture, we fit a plane through the origin to the set 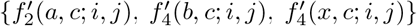 with (*k* − 4)(*k* − 5)/2 points.

This finishes the instrumental part. Important remarks should also be made in addition. (a) Two different rationales are the basis of the geometric and the standard framework derivations and the approximate solutions also differ formally. In 2-way admixture simulations, both estimates are quite similar. (b) The introgression regime is more quickly affected by post-admixture drift. The time to fool the admixture test decreases linearly with the proportion of the introgressor, as compared to an even mixture. (c) Auxiliary populations having large shared drift with donors improve the accuracy of proportion estimates. (d) Auxiliary populations with small shared drift should not impair the estimates because they accumulate at the graphical representation origin. However, they can give unrealistic determinations in the direct computation of the *f*_4_ ratio. These observations may be relevant in unsupervised computations.

The idea of using a graphical visualization of the population admixture problem may help us better grasp it and foster new applications. In this respect, we have introduced the angles *ϕ* and φ, pre and post JL projection, respectively. Whereas the value of *ϕ* is only a primary indicator to witness a population admixture, the value of φ is significant in assessing the reconstruction quality of the native admixture configuration. Also, on the basis of the Brownian trajectories that populations may have taken in phase space, we may also understand the amplified impact that genetic drift has on introgression in comparison to an even mixture. Eventually, we cite an interesting *rule of thumb* proposed in [7] in connection with the interpretation of PCA plots, commonly used to study population structure, where the own geometric interpretation of the constraint (16), cos *ϕ* < 0, leads to the approximate recipe that in such PCA plots, given two admixture contributors then the putative hybrids should be inside the circle whose diameter is determined by the two donors.

We end this discussion with the reminder that the population admixture problem has facets other than the postadmixture drift that are less tractable. The geometric view provides fresh perspectives to the essentially statistical foundations of the *f*-formalism that could help us better grasp untackled aspects.

## Acknowledgements

JAO work has been partially funded by the project PID2019-109592GB-I00/AEI/10.13039/501100011033 from the Spanish Ministerio de Ciencia e Innovación Agencia Estatal de Investigación.

## Appendix

Some features of the tetrahedron in terms of edge lengths are gathered. Let V be the tetrahedron volume, A the base area, and *h* the height (determined by the hybrid vertex). We have V = Ah/3.

The length of edge 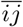is d_*ij*_ and we assume that the hybrid is located at vertex 1 The volumen may be calculated using the six edge lengths with the Cayley-Menger determinant [20, 18]

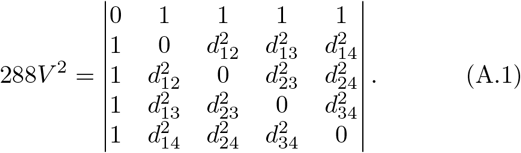

In the allele frequency space, 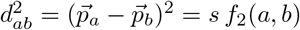. In the JL subspace, 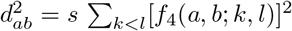 Explicitly

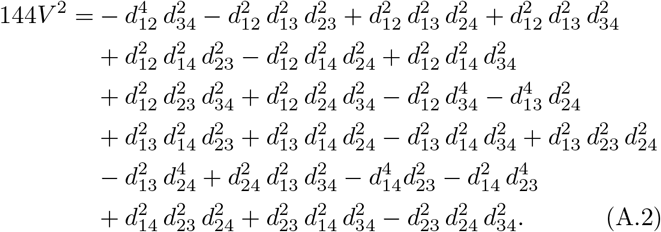

The base area may be computed with the Heron formula [18], which involves the three triangle edges

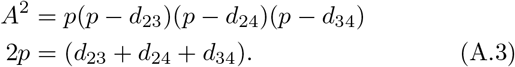

Combining those results we get the tetrahedron height, *h* = 3V/A, in terms of its edge lengths and define a dimensionless flatness index as the ratio 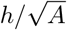. In terms of volume and base area it reads: 3V/A^3*/*2^, to be computed with (A.2 and (A.3). The Cayley-Menger determinant (A.1) can be used to show explicitly that in a 3-way admixture at birth the four populations form a co-planar configuration. It suffices to use the linear combination 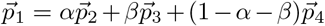, to compute the distances d_1,*k*_, with *k* = 2, 3, 4, in (A.1). The result of the determinant factorizes as 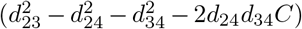, where Q is a factor that is irrelevant for our goal and, more crucially, C is the cosine of the angle between edges 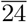 and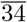. The determinant vanishes because the first factor does as a result of the cosine law.

## Footnotes

1 In the fit *y* = *m x* to a set of *n* points, the slope reads 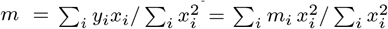 where *m*_*i*_ = *y*_*i*_*/x*_*i*_ is the slope defined by the point (*x*_*i*_, *y*_*i*_), and *i* = 1, …, *n*.

## References

[1] Nick Patterson, Priya Moorjani, Yontao Luo, Swapan Mallick, Nadin Rohland, Yiping Zhan, Teri Genschoreck, Teresa Webster, and David Reich. Ancient admixture in human history. Genetics, 192:1065–93, 2012.

[2] Wolfgang Haak, Iosif Lazaridis, Nick Patterson, Nadin Rohland, Swapan Mallick, Bastien Llamas, Guido Brandt, Susanne Nordenfelt, Eadaoin Harney, Kristin Stewardson, Qiaomei Fu, Alissa Mittnik, Eszter Bánffy, Christos Economou, Michael Francken, Susanne Friederich, Rafael Garrido Pena, Fredrik Hallgren, Valery Khartanovich, Aleksandr Khokhlov, Michael Kunst, Pavel Kuznetsov, Harald Meller, Oleg Mochalov, Vayacheslav Moiseyev, Nicole Nicklisch, Sandra L. Pichler, Roberto Risch, Manuel A. Rojo Guerra, Christina Roth, Anna Szécsényi-Nagy, Joachim Wahl, Matthias Meyer, Johannes Krause, Dorcas Brown, David Anthony, Alan Cooper, Kurt Werner Alt, and David Reich. Massive migration from the steppe was a source for Indo-European languages in Europe. Nature, 522:207–211, 2015.

[3] Iosif Lazaridis, Dani Nadel, Gary Rollefson, Deborah C. Merrett, Nadin Rohland, Swapan Mallick, Daniel Fernandes, Mario Novak, Beatriz Gamarra, Kendra Sirak, Sarah Connell, Kristin Stewardson, Eadaoin Harney, Qiaomei Fu, Gloria Gonzalez-Fortes, Eppie R. Jones, Songül Alpaslan Roodenberg, György Lengyel, Fanny Bocquentin, Boris Gasparian, Janet M. Monge, Michael Gregg, Vered Eshed, Ahuva Sivan Mizrahi, Christopher Meiklejohn, Fokke Gerritsen, Luminita Bejenaru, Matthias Blüher, Archie Campbell, Gianpiero Cavalleri, David Comas, Philippe Froguel, Edmund Gilbert, Shona M. Kerr, Peter Kovacs, Johannes Krause, Darren McGettigan, Michael Merrigan, D. Andrew Merriwether, Seamus O’Reilly, Martin B. Richards, Ornella Semino, Michel Shamoon-Pour, Gheorghe Stefanescu, Michael Stumvoll, Anke Tönjes, Antonio Torroni, James F. Wilson, Loic Yengo, Nelli A. Hovhannisyan, Nick Patterson, Ron Pinhasi, and David Reich. Genomic insights into the origin of farming in the ancient Near East. Nature, 536:419–424, 2016.

[4] Benjamin M. Peter. Admixture, Population Structure, and F-Statistics. Genetics, 202:1485–501, 2016.

[5] Mark Lipson. Applying f4-statistics and admixture graphs: Theory and examples. Molecular Ecology Resources, 20:1658–1667, 2020.

[6] Eádaoin Harney, Nick Patterson, David Reich, and John Wakeley. Assessing the performance of qpAdm: A statistical tool for studying population admixture. Genetics, 217:iyaa045, 2021.

[7] Benjamin M. Peter. A geometric relationship of *f*_2_, *f*_3_ and *f*_4_-statistics with principal component analysis. Philosophical Transactions of the Royal Society B: Biological Sciences, 377:20200413, 2022.

[8] L.L. Cavalli-Sforza. Population structure and human evolution. Proceedings of the Royal Society of London. Series B, 164:362–379, 1966.

[9] L.L. Cavalli-Sforza and A. Piazza. Analysis of evolution: Evolutionary rates, independence and treeness. Theoretical Population Biology, 8:127–165, 1975.

[10] L.L. Cavalli-Sforza, P. Menozzi, and A. Piazza. The history and geography of human genes. Princeton University Press, 1994.

[11] J. C. Long. The genetic structure of admixed populations. Genetics, 127:417–428, 2 1991.

[12] Gonzalo Oteo-García. Archaeogenetics of Southwest Europe. http://eprints.hud.ac.uk/id/eprint/35459/, xPhD thesis, 2020.

[13] Lily Agranat-Tamir, Shamam Waldman, Naomi Rosen, Benjamin Yakir, Shai Carmi, and Liran Carmel. Linadmix: evaluating the effect of ancient admixture events on modern populations. Bioinformatics, 37:4744–4755, 2021.

[14] Gonzalo Oteo-García and José-Angel Oteo. A Geometrical Framework for f-Statistics. Bulletin of Mathematical Biology, 83:14, 2021.

[15] Roman Vershynin. High-Dimensional Probability: An Introduction with Applications in Data Science. Cambridge University Press, 2018.

[16] Mark Lipson, Po-Ru Loh, Alex Levin, David Reich, Nick Patterson, and Bonnie Berger. Efficient Moment-Based Inference of Admixture Parameters and Sources of Gene Flow. Molecular Biology and Evolution, 30(8):1788–1802, 05 2013.

[17] David Reich Lab. Github-DReichLab/AdmixTools: Tools test whether admixture occurred and more. https://github.com/DReichLab/AdmixTools, 2023.

[18] Duncan M. Y. Sommerville. Introduction to the Geometry of N Dimensions. Courier Dover Publications, 2020.

[19] Swapan Mallick and David Reich. The Allen Ancient DNA Resource (AADR): A curated compendium of ancient human genomes. 10.7910/DVN/FFIDCW, 2023. Harvard Dataverse, V1.

[20] Karl Wirth and André S. Dreiding. Edge lengths determining tetrahedrons. Elemente der Mathematik, 64:160–170, 2009.

